# Visual predictions, neural oscillations and Newtonian physics

**DOI:** 10.1101/2020.02.19.951566

**Authors:** Blake W. Saurels, Wiremu Hohaia, Kielan Yarrow, Alan Johnston, Derek H. Arnold

**Affiliations:** School of Psychology, The University of Queensland, Australia; Department of Psychology, City, University of London, United Kingdom; School of Psychology, University of Nottingham, United Kingdom

**Keywords:** Visual Predictions, Neural Oscillations, Naturalistic Visual Input

## Abstract

Prediction is considered a core function of the human visual brain, but relating this suggestion to real life is problematic, as findings regarding the neural correlates of prediction rely on abstracted experiments, not reminiscent of a typical visual diet. We addressed this by having people view videos of basketball, and asking them to predict jump shot outcomes while we recorded eye movements and brain activity. We used the brain’s understanding of physics to manipulate predictive success, by inverting footage. People had enhanced alpha-band activity in occipital brain regions when watching upright videos, and this predicted both an increase in predictive success and enhanced ball tracking. Alpha-band activity in visual brain regions has been linked to inhibition, so we regard our results as evidence that inhibition of task irrelevant information is a core function of predictive processes in the visual brain, enacted as people complete visual tasks typical of daily life.

---

In daily life, people are exposed to an abundance of sensory information at any given moment. The human brain does not process all this information equally - instead, it implements filtering processes, selecting only a subset of input for detailed analyses [1-4]. One property that can determine if an input will be selected for detailed analysis is if it is surprising [5]. Studies using tightly controlled inputs, not reminiscent of our daily visual diet (e.g., trains of flashed stimuli), suggest the human brain is more responsive to inputs that have *not* been predicted [6, 7].

While there is great interest in the processes underlying prediction, of both visual events and the activity they trigger [e.g., 8, 9], the highly abstract and controlled nature of stimuli used in most prediction experiments limit the degree to which results can speak to neural processing in daily life. We address this issue by examining how well people can predict outcomes while watching inputs that are more reminiscent of daily life – videos of basketball, culminating in a jump shot (see Figure 1g). In addition to asking participants to predict the shot outcome, we ask them to track the ball as it moves. We manipulate peoples’ ability to predict by presenting videos upright, a situation that accords with peoples’ intuitive understanding of physics [10, 11], or upside-down, a situation that conflicts with intuitive expectations. We track peoples’ eye movements and take electroencephalogram (EEG) recordings. Of particular interest is alpha-band (8-12Hz) oscillatory activity from visual brain regions, as these have been implicated as a neural marker of inhibited information processing during selective attention, originating in deep layers of visual cortex [12-16]. Similar processes might underlie the suppression of information processing when events can be anticipated. Thus, we predicted that, for upright relative to inverted videos, people would be more accurate at predicting shot outcomes and tracking the ball position, and would have enhanced occipital alpha-band oscillatory activity.

**Figure 1.**
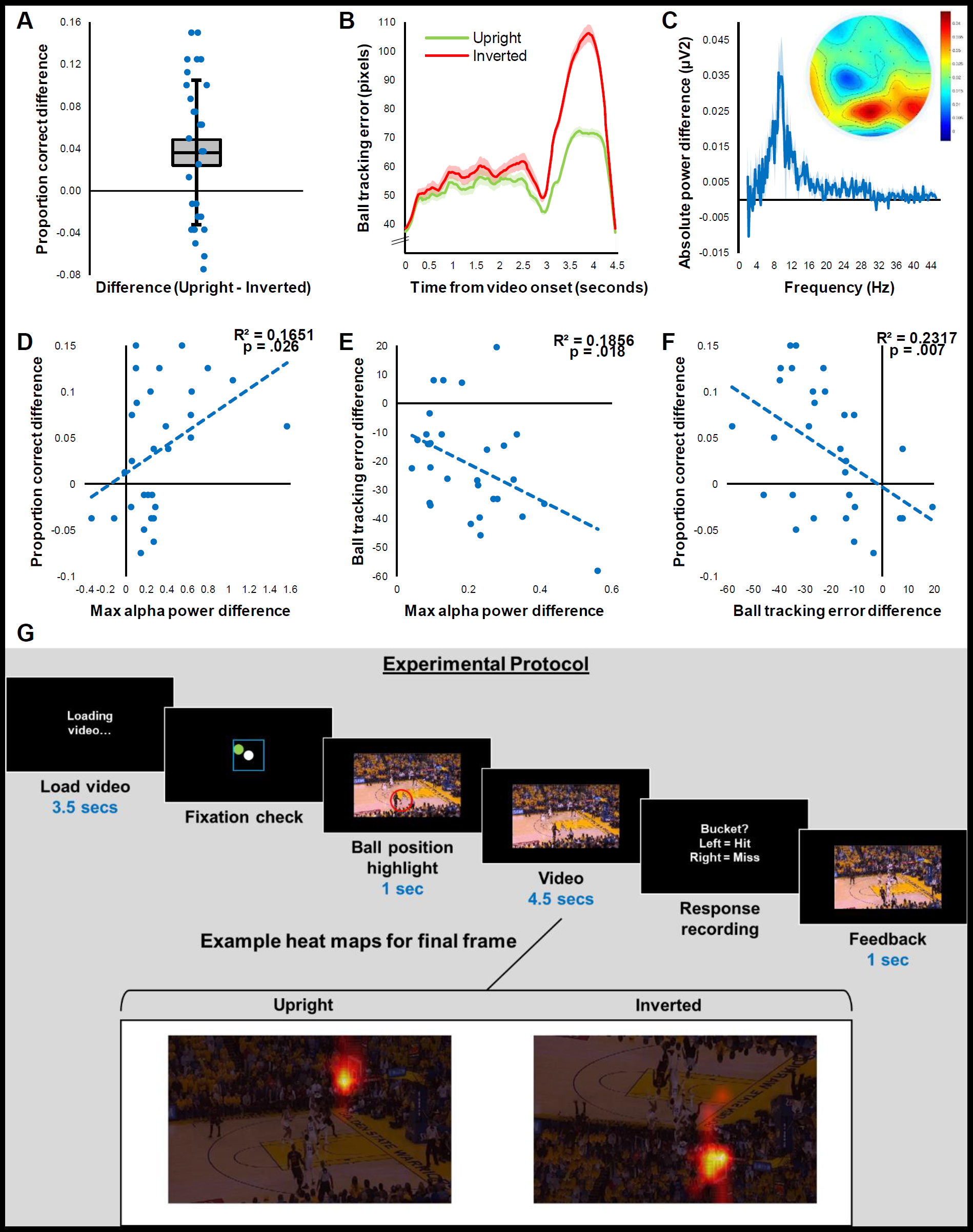
**A)** Proportion correct difference scores (blue) for individual participants. Central black line depicts median score. Light grey shaded region depicts ±1 standard error. Black error bar depicts ±1 standard deviation. Coloured circles depict individual scores. **B)** Mean error (pixels), between individual fixation positions and actual ball locations, as a function of time from video onset (seconds) for upright (green) and inverted (red) videos. Shaded regions depict SE across participants. **C)** Absolute power difference (*µV*^2^) as a function of frequency (Hz). Shaded regions depict SE across participants. Topographic map depicts conditional difference in alpha-band (8-12Hz) activity for full sample duration. **D)** Conditional difference in proportion correct as a function of difference in max absolute alpha power (*µV*^*2*^) – with spectra analysis restricted to the last 250ms of each video. **E)** Conditional difference in ball tracking error as a function of difference in max absolute alpha power (*µV*^*2*^) – with analysis restricted to the last 1.5 seconds of each video. **F)** Conditional difference in proportion correct as a function of difference in ball tracking error – with analysis restricted to the last 1.5 seconds of each video. **G)** Graphic depicting experimental protocol and example heat maps for final frame of a video.

## RESULTS

As expected, we found that overall people were *better* at predicting outcomes when footage was upright, as opposed to inverted (*t*_29_ = 2.8856, *p* = .007, *BF*_10_ = 5.884; see Figure 1a). We also found that participants were better at tracking the ball as it moved in upright, as opposed to inverted, videos (*t*_29_ = 10.495, *p* < .001, *BF*_10_ = 4.243e+8; see Figure 1b), and that alpha-band oscillatory activity was enhanced while watching upright, as opposed to inverted, videos (*t*_29_ = 4.349, *p* < .001, *BF*_10_ = 176.462; see Figure 1c).

Conditional differences in alpha-band activity (upright minus inverted), averaged across the full video sample, did not predict conditional differences in predictive accuracy (*R*^*2*^ = .0015, *p* = .838, *BF*_10_ = 0.232). We reasoned this might be due to early epochs being less relevant for the prediction of events at the end of the video. So, we restricted a repeated analysis to oscillatory activity *immediately prior* to the jump shot [as in previous similar investigations, see 12] – the final 250ms of each video. This revealed a *positive* relationship between conditional differences in alpha-band activity and predictive accuracy (*R*^*2*^ = .1651, *p* = .026, *BF*_10_ = 2.426; see Figure 1d).

Conditional differences in ball tracking error were found primarily for the last 1.5 seconds of video footage (see Figure 1b) – a period associated with enhanced ball movement and the jump shot itself. Looking at this portion of the video sample, conditional differences in ball tracking error were *negatively* associated with both occipital alpha-band activity (*R*^*2*^ = .1856, *p* = .018, *BF*_10_ = 3.361; see Figure 1e), and predictive accuracy (*R*^*2*^ = .2316, *p* = .007, *BF*_10_ = 7.216; see Figure 1f).

Our instruction to track the ball while watching videos raises the possibility that saccadic activity might have differed between the two conditions, causing people to make more or less predictive errors, and disrupting alpha-band activity. We found that participants did make fewer saccades while watching upright videos (*t*_29_ = 5.663, *p* < .001, *BF*_10_ = 5294.371). We therefore conducted two hierarchical multiple regressions (HMR; see Methods for analysis details) to determine if saccades might underlie the other relationships we have uncovered. Saccade numbers *did not* predict predictive accuracy (*R*^*2*^ *Ch*. = .001, *F Ch*._*28*_ = 0.022, *p* = .882). Alpha-band activity (*R*^*2*^ *Ch*. = .176, *F Ch*._*27*_ = 5.785, *p* = .023) and ball tracking error (*R*^*2*^ *Ch*. = .241, *F Ch*._*27*_ = 8.563, *p* = .007) *both independently* explained additional variance in predictive accuracy, after accounting for the influence of saccade numbers – showing that neither of these effects could be ascribed to saccade number differences.

One possible criticism of our findings is that the power of alpha oscillations might have scaled with the ease of the task, rather than these being indicative of a difference in underlying neural processes. To assess this, we identified a subset of 30 videos on which participant performance had been *matched* across conditions (*t*_29_ = 0.682, *p* = .501, *BF*_10_ = 0.241). The alpha difference observed before shot outcomes using *all* clips (*t*_29_ = 2.828, *p* = .008, *BF*_10_ = 5.214) persisted for the sub-set of trials for which performance was matched (*t*_29_ = 3.626, *p* = .001, *BF*_10_ = 30.654; see Figure 2).

**Figure 2.**
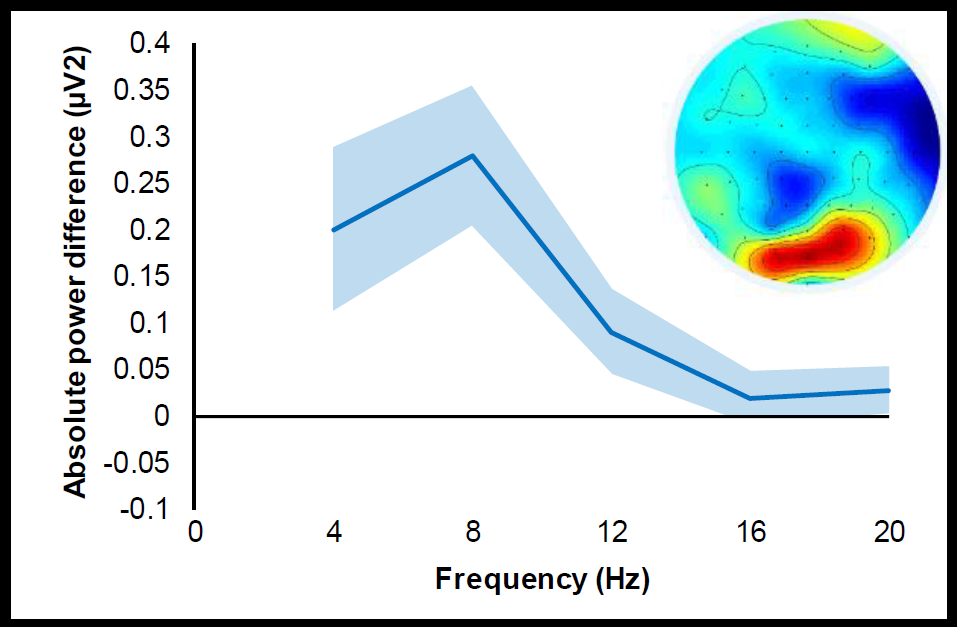
Absolute power difference (*µV*^2^) in occipital activity for the last 250ms of each video, as a function of frequency (Hz) - using a subset of videos where performance was matched across conditions. Shaded regions depict SE across participants. Topographic map depicts conditional difference in alpha-band (8-12Hz) activity.

## DISCUSSION

We find that for more predictable, upright videos, people: were better at predicting shot outcomes, were better at tracking the ball, and had enhanced occipital alpha-band oscillatory activity. These measures were also intercorrelated, such that greater alpha-band activity immediately before taking a shot predicted a better predictive outcome and more accurate ball tracking, and that more accurate ball tracking was associated with better outcome prediction.

Differences in alpha-band activity have previously been ascribed to fatigue, which can be *positively* related with alpha-band activity [17, 18]. We think it is unlikely that this could account for our data, as we have found that alpha-band activity is associated with *better*, not worse performance. It could also be suggested that one of our core findings, that the power of alpha oscillations is enhanced for upright relative to inverted video presentations, is indicative of differences in task difficulty, as opposed to the recruitment of distinct neural processes. However, we can discount this possibility as when we identified a subset of videos that had resulted in matched levels of performance across upright and inverted video presentations, we found that the power of alpha oscillations was still enhanced for upright relative to inverted video presentations. This shows that these conditions had not just encouraged different levels of performance, but rather that they had additionally recruited separate neural processes, with distinguishable neural signatures.

Enhanced alpha-band (8-12Hz) oscillatory activity is a well-established neural signature of spatial-inhibition in the visual brain [12-16]. We therefore regard our results as evidence for the implementation of a dynamic form of spatial attention, guided by visual predictions. As people are attuned to dynamics within an environment with gravity, they were better able to predict shot trajectories when footage was upright, and to track balls as they moved (see Figure 1). We believe these behavioural signatures of enhanced visual prediction were coupled with a neural signature – an increase in alpha-band activity across visual brain regions, as the processing of input from positions *other* than those predicted to be relevant was suppressed [see 13; i.e., not around the ball].

Overall, we have found that when people attempt to predict the outcome of basketball jump shots, they perform better when input conforms to gravity, and that this encourages a neural signature characterised by a greater power of alpha oscillations across visual brain regions. Moreover, this neural signature is indicative of different neural processes, as it persists even if analyses are restricted to inputs that encourage matched levels of performance.

## METHOD

### Experiment

#### Participants

Thirty participants via a first-year research participation scheme at the University of Queensland. The experiment was restricted to participants with normal vision (glasses and contact lenses interfered with the eye tracker). Ages ranged from 18 to 29 (*M* = 20.53, *SD* = 2.92).

#### Stimuli and apparatus

Eighty 5 second basketball clips were encoded from two full basketball games. These all contained a jump shot where the ball reached the basket at the 137th frame. These had a width subtending 20.04 degrees of visual angle (dva), and a height subtending 9.94 dva. Videos were presented on an ASUS VG248QE 3D Monitor, using the Psychophysics Toolbox software through MatLab R2015B, at 30 frames per second. The monitor had a resolution of 1920 × 1080 pixels and was set at a refresh rate of 60Hz (updated twice per video frame). Participants viewed stimuli at a distance of 57cm, from directly in front of the monitor with their chin placed on a chin rest. A Cambridge Research Systems LiveTrack 60 Hz fixation monitor was used for eye tracking. A Biosemi International ActiveTwo system was used to record EEG data (sampling rate: 1024Hz, channel array: 64).

#### Procedure

Each participant watched all 80 clips twice, once upright and once inverted. Presentation order was counter-balanced such that 40 clips were seen upright first, and 40 inverted first, so any conditional differences could not be attributed to practice or learning. This also allowed for each clip to serve as its own control.

At the start of the experiment, participants performed a nine-point eye tracking calibration. A short test of the eye tracking calibration (a moving white dot was presented for the participant to follow, with eye gaze data presented onscreen) and a practice trial was then completed. These were used to confirm the calibration quality, and to explain the task.

At the start of each trial, participants performed a modified fixation check. A white fixation dot (0.44 dva) was shown in the centre of the screen, and a green dot (0.48 dva) marked the eye gaze position as registered by the eye tracker. If the participant was looking at the fixation dot, and the eye tracking dot was on or within 35 pixels (0.97 dva; marked by a blue square) of the fixation dot, then they could initiate the trial with the middle mouse button. If they were fixating and the eye tracking dot was offset, this would indicate to the participant that they had altered their head position and would need to restore it to the initial calibration position. This insured the eye tracking calibration quality was maintained throughout the experiment.

Directly after the fixation check, the first frame of the video was presented for 1 second. An open, red, shrinking circle (2.48 dva to 0.83 dva) was overlaid on the ball position for this period. After this, 4.5 seconds (135 frames) of basketball footage was played, generally depicting some dribbling and passing, and culminating in a jump shot. The final frame depicted the ball position 2 frames before reaching the basket (or backboard; the frame in which you could tell if the ball would go in or miss). The screen then displayed two mouse options concerning the behavioural judgement: left mouse if they thought it would be a bucket, right if they thought it would be a miss. The participant then received feedback on their response; the last 1 second of the video was shown, and the video edge was outlined with either a green (correct) or red (incorrect) rectangle. An experiment progression circle was then briefly flashed, and then the next video would begin loading, which took roughly 3-4 seconds.

Behavioural responses, trial order, and eye tracking data were recorded in a series of MatLab structures that were saved periodically throughout the experiment into .mat files.

### Data Analysis

#### Behavioural data

The proportion of trials individual participants correctly reported the shot outcome was compared for upright and inverted videos, using paired sample frequentist and Bayesian *t*-tests (*null*: no conditional difference, *alternative*: conditional difference present).

#### Ball tracking data

Eye gaze position, taken as screen pixel position relative to screen centre, was recorded for each frame of the videos. The left eye was used primarily, except in cases where calibration proved difficult with the left, and so the right eye was used instead.

For each frame of each video, the error between individual fixation positions and actual ball location was calculated (as a vector magnitude). Paired sample frequentist and Bayesian *t*-tests were used to compare each participant’s mean error scores (pixels) for upright and inverted trials (*null*: no conditional difference, *alternative*: conditional difference present).

#### Frequency data

During pre-processing, data were high (1Hz), low (100Hz) and band-pass (45 – 55Hz) filtered. Data were then subjected to an independent component analysis, implemented by the FieldTrip toolbox for Matlab, to remove blink artefacts. Electrode activity was then average referenced, to correct for baseline skin conductance levels. After pre-processing, trial sorting and spectra analyses were conducted, via custom Matlab scripts using the fast Fourier transform.

Data presented in Figure 1C is averaged across all channels. The topographical map highlights occipital involvement, as expected.

Mean individual alpha-band oscillatory activity was compared for upright and inverted trials using paired sample frequentist and Bayesian *t*-tests (*null*: no conditional difference, *alternative*: conditional difference present).

#### Saccades

The number of saccades performed by each participant on each trial was calculated using custom MatLab code. An eye movement was classified as a saccade if, for a period greater than 2 frames (∼66ms), the gaze change velocity exceeded 5 pixels per frame (0.14 dva/∼33ms) and the angle of movement varied by less than 30° relative to the previous frames’ angle of movement. Multiple examples of this rudimentary saccade classifier can be found in the data repository. Paired sample frequentist and Bayesian *t*-tests were used to compare each participant’s mean number of saccades per trial for upright and inverted videos (*null*: no conditional difference, *alternative*: conditional difference present).

#### Correlations

Individual conditional differences were calculated to examine the relationship between the three key data points recorded: proportion correct, ball tracking error, and alpha-band oscillatory activity. These differences were always taken as upright minus inverted trials. The (Bayesian) Pearson correlations between these three sets of difference scores were calculated (*null*: no relationship, *alternative*: relationship present).

For the ball tracking data, difference scores were calculated using data from the last 1.5 seconds of each video, as this period contained the most physical ball movement (e.g., the jump shot). When this data was correlated with alpha-band oscillatory difference scores, a matched period of neural activity was examined. When correlating alpha-band oscillatory difference scores with proportion correct difference scores, a short period *immediately prior* to the behavioural judgement was used (250ms; see 10). Alpha-band oscillatory scores were defined as the max power within the 8-12Hz range, taken from electrodes: PO7, PO3, O1, Oz, POz, PO8, PO4, and O2 – a pre-determined occipital cluster.

#### Hierarchical multiple regressions

Two HMRs were conducted to determine the influence of saccadic activity on the predictive relationships that explained behaviour (conditional differences in predictive accuracy). Accordingly, individual conditional differences in mean saccades per trial were entered at Step 1 in both regressions. This did not predict variance in predictive accuracy (*R*^*2*^ *Ch*. = .001, *F Ch*._*28*_ = 0.022, *p* = .882). Conditional differences in ball tracking error and alpha-band activity were entered at Steps 2 and 3, in both orders. When alpha-band activity was entered at Step 2, additional variance in predictive accuracy was explained over Step 1 (*R*^*2*^ *Ch*. = .176, *F Ch*._*27*_ = 5.785, *p* = .023), and when ball tracking error was entered at Step 3, additional variance in predictive accuracy was explained over Step 2 (*R*^*2*^ *Ch*. = .134, *F Ch*._*27*_ = 5.076, *p* = .033). Alternatively, when ball tracking error was entered at Step 2, additional variance was explained over Step 1 (*R*^*2*^ *Ch*. = .241, *F Ch*._*27*_ = 8.563, *p* = .007), but when alpha-band activity was entered at Step 3, no additional variance was explained over Step 2 (*R*^*2*^ *Ch*. = .070, *F Ch*._*27*_ = 2.649, *p* = .116). This suggests that all the variance in predictive accuracy explained by alpha-band activity is also captured by shared variance explained by ball tracking error.

#### Task difficult matching

To control for the possibility that task difficult solely explained the conditional differences in alpha-band occipital activity observed, a subset of the videos was selected on which shot outcome performance was matched. Videos pairs (upright & inverted) with a difference in proportion correct (across participants) of less than or equal to 0.05 where isolated. Thirty clips met this criteria, and the average proportion correct for each condition was 0.88. The spectra differences presented in Figure 2 were computed using the same occipital cluster as above, for the performance-matched subset.

